# Intracellular mechanics and TBX3 expression jointly dictate the spreading mode of melanoma cells in 3D environments

**DOI:** 10.1101/2022.06.09.495509

**Authors:** Ghodeejah Higgins, Faatiemah Higgins, Jade Peres, Dirk M Lang, Tamer Abdalrahman, Muhammad H. Zaman, Sharon Prince, Thomas Franz

**Affiliations:** Biomedical Engineering Research Centre, Division of Biomedical Engineering, Department of Human Biology, University of Cape Town, Observatory, South Africa; Division of Cell Biology, Department of Human Biology, University of Cape Town, Observatory, South Africa; Department of Biomedical Engineering, Boston University, Boston, MA, USA; Howard Hughes Medical Institute, Chevy Chase, MD, USA; Bioengineering Science Research Group, Faculty of Engineering and Physical Sciences, University of Southampton, Southampton, UK

**Author notes:** Corresponding author, Department of Human Biology, Faculty of Health Sciences, University of Cape Town, Private Bag X3, Observatory 7935, South Africa.

**Keywords:** Cluster formation, cluster morphology, mitochondrial particle tracking microrheology, cell mechanics, T-box factor, collagen

## Abstract

Cell stiffness and T-box transcription factor 3 (TBX3) expression have been identified as biomarkers of melanoma metastasis in 2D environments. This study aimed to determine how mechanical and biochemical properties of melanoma cells change during cluster formation in 3D environments. Vertical growth phase (VGP) and metastatic (MET) melanoma cells were embedded in 3D collagen matrices of 2 and 4 mg/ml collagen concentrations, representing low and high matrix stiffness. Mitochondrial fluctuation, intracellular stiffness, and TBX3 expression were quantified before and during cluster formation. In isolated cells, mitochondrial fluctuation decreased and intracellular stiffness increased with increase in disease stage from VGP to MET and increased matrix stiffness. TBX3 was highly expressed in soft matrices but diminished in stiff matrices for VGP and MET cells. Cluster formation of VGP cells was excessive in soft matrices but limited in stiff matrices, whereas for MET cells it was limited in soft and stiff matrices. In soft matrices, VGP cells did not change the intracellular properties, whereas MET cells exhibited increased mitochondrial fluctuation and decreased TBX3 expression. In stiff matrices, mitochondrial fluctuation and TBX3 expression increased in VGP and MET, and intracellular stiffness increased in VGP but decreased in MET cells. The findings suggest that soft extracellular environments are more favourable for tumour growth, and high TBX3 levels mediate collective cell migration and tumour growth in the earlier VGP disease stage but play a lesser role in the later metastatic stage of melanoma.

**Symbols:** 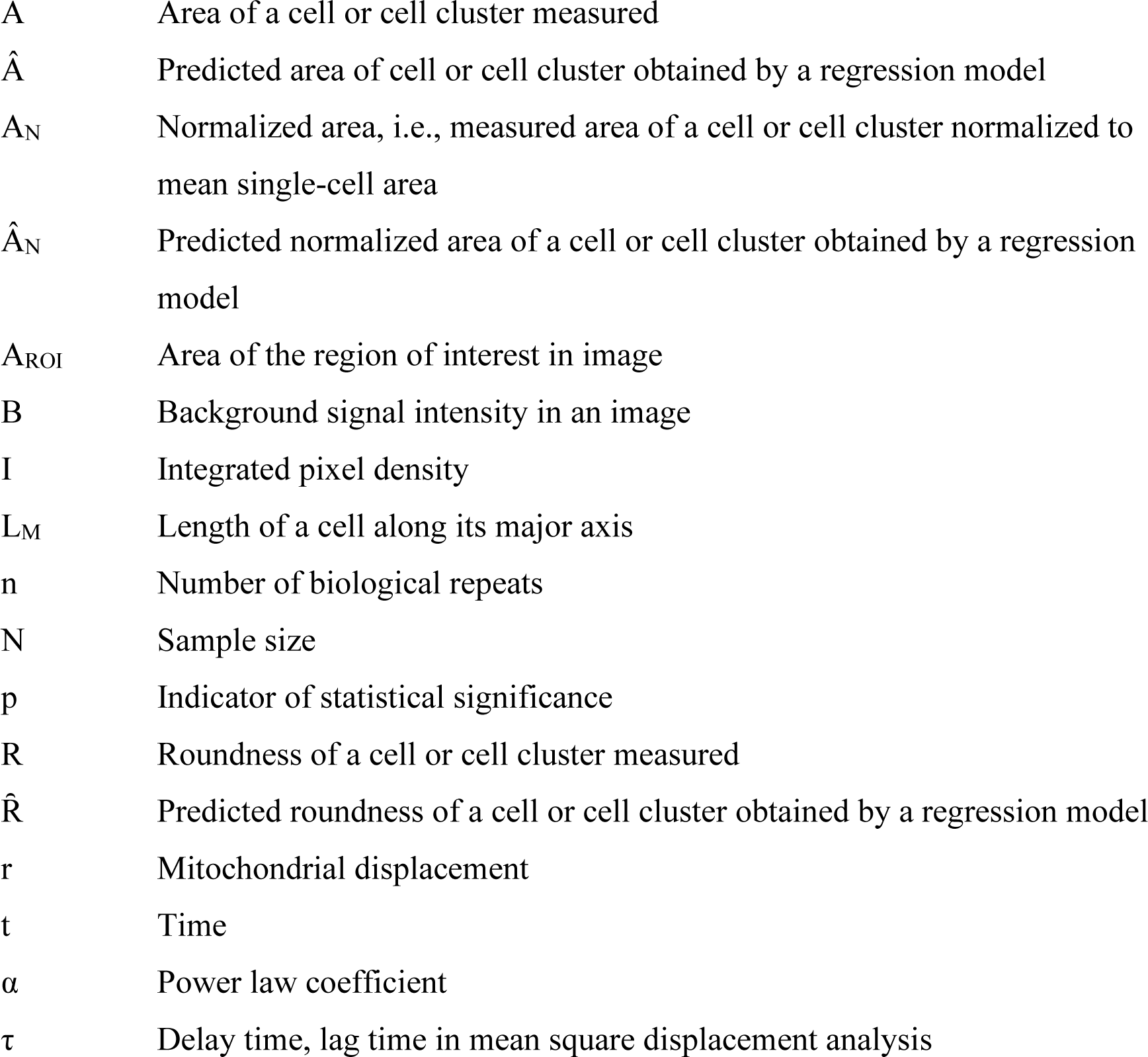

## 1. Introduction

Recent studies have shown that cancer cells continuously respond and adapt to changes in the microenvironment through altered migratory properties [1], mechanical properties [2], and gene expression [3, 4]. Some studies focused on the mechanical properties of cells in isolation, whereas others investigated the mechanical properties of cellular aggregates. For cells in isolation, it was shown that cancer cells have enhanced biomechanical properties compared to normal cells, such as increased contractile [5] and traction force generation [1, 6, 7] and deformability [1, 8, 9]. At the cellular aggregate level, compromised rigidity of solid cancer tumours compared to normal tissue was demonstrated by measuring stiffness averaged over thousands of cells in cell spheroid models using many techniques, such as micro-tweezers [10], atomic force microscopy (AFM) [11], and cavitation rheology [12].

These studies have provided insight into the cell mechanics of various cancer types and identified mechanical properties as potential biomarkers of cancer malignancy; however, their results have not demonstrated the progressive changes in biomechanical properties as cells transition from isolated single cells to cellular aggregates. In particular, it is not understood whether isolated cells from different stages of the disease are innately endowed with sufficient mechanical properties to form tumours or if mechanical properties are adjusted in response to three-dimensional (3D) extracellular matrix (ECM) properties at distant sites of colonization.

It is increasingly being recognized that the properties of the tumour microenvironment play an essential role in cancer development [13]. Cells are surrounded by a rich ECM, an essential component in the microenvironment, whose chemical, physical, and mechanical properties are fundamental in orchestrating several biological processes, including tissue homeostasis, differentiation, migration and invasion [14]. The most abundant protein found in the ECM is fibrillar type I collagen [15, 16], which forms a scaffold that imparts tensile strength and stiffness to the tissue [17, 18]. It has also been reported that ECM stiffness changes are due to altered collagen I density [18-21]. In addition, collagen I comprises 80-90% of the collagenase proteins in the dermis, and studies have shown that altered collagen I density affects melanoma progression *in vitro* and *in vivo* [22].

Cell-ECM interaction is mediated through mechanical machineries such as the cell’s actin cytoskeleton, molecular motors, integrins, focal adhesion and GTPases [23-25], allowing cells to receive, process, and respond to biochemical and biomechanical cues from the environment [26, 27]. The properties, including composition, topography, and stiffness of tumour-derived ECM, are enhanced compared to normal ECM and crucial in administering cells with a tumorigenic genotype [19].

Although ECM mechanics and its role in guiding metastasis have received increasing attention [15, 28-31], little is known about the biomechanics of the cell-ECM relationship at distant sites of metastasis and why cells may favour specific distant organs over others. Even less is known about the cell’s inherent biomechanical properties and biochemical pathways that coordinate formation of tumours in changing 3D environments, particularly at advanced stages of cancer progression. The ability of cells to form clusters is essential for tumour formation and metastasis [32].

Several assays, together with high-resolution imaging approaches, have addressed essential questions on cluster morphology, tumour growth, viability [33, 34], drug testing [4] and drug development [33, 35, 36], spheroid invasion [37-39] and cancer stem cell content [4]. Fewer studies have investigated the contribution of cell mechanics in the context of cell deformability to the detachment and invasion process. The scarcity of cell mechanics studies using cell clusters and spheroids is partly due to technical challenges in quantifying cell mechanics in 3D environments, particularly the detachment and invasion process in a live-cell 3D realistic model. Microrheological methods, particularly those facilitating the tracking of endogenous nano-particles such as mitochondria inside the cells, can offer non-invasive techniques to passively probe cell mechanics without introducing cellular or matrix deformation [2]. Furthermore, local responses are probed, making it convenient to investigate cell mechanics in complex environments where single and collective cell behaviour may co-exist.

A central therapeutic goal has been to target key molecular regulators of advanced-stage cancers. The AKT3 signalling pathway plays a critical role in melanoma formation and invasion, and we have reported that TBX3 is a key substrate of AKT3 in melanomagenesis [40]. We have shown that AKT3 phosphorylates TBX3 at S720, promoting TBX3 protein stability, nuclear localization, transcriptional repression of E-cadherin, and its role in cell migration and invasion. We have also previously reported that TBX3 levels are high in VGP and metastatic melanoma and that depleting TBX3 in these cells inhibits their ability to form tumours [41]. Consistent with these findings, when TBX3 alone was highly expressed in RGP non-malignant melanoma cells, it was sufficient to promote tumour formation and invasion [42].

Although it is known that TBX3 is highly expressed in several cancers [43], knowledge of mechanical mediators of TBX3 regulation, such as matrix rigidity, is scarce. Research on TBX3 has been conducted predominantly in 2D *in vitro* conditions and in *in vivo* animal models. However, cells in 2D environments do not fully depict physiologically cell-cell and cell-matrix interactions [44]. *In vivo* models mimic complex cell-cell and cell-matrix interaction better but exhibit spatially complex microenvironments whose environmental cues, as ECM properties, are challenging to control [45].

Considering that TBX3 promotes tumour formation in melanoma and tumour formation is dependent on the mechanical environment, the current study aimed to determine how mechanical mediators such as matrix rigidity and cell stiffness impact (i) cluster formation of advanced melanoma cells at the vertical growth phase and metastatic stage and (b) TBX3 expression in these melanoma cells during cluster formation in 3D *in vitro* environments.

## 2. Materials and methods

### 2.1. Cell seeding

Advanced human melanoma cell lines, ME1402 vertical growth phase (VGP) and WM1158 metastatic (MET), were cultured using RPMI 1640 media (Life Technologies), supplemented with 10% foetal bovine serum (FBS) and 1% penicillin-streptomycin. Before passaging or experimentation, cells were grown to near-confluence in 25 cm^2^ tissue flasks (Costar, Corning Life Science, Acton, MA) at 37°C and 5% CO_2_. Media was replaced every 3-4 days.

### 2.2. Collagen embedding

Single cells were embedded in 2 and 4 mg/ml collagen solutions. Collagen solutions comprised Type I Rat Tail Collagen (BD Biosciences, stock solution of 8.21 mg/ml) and neutralizing solution (100 mM HEPES in 2X PBS with pH 7.3) in a 1:1 ratio, diluted with 1XPBS to final concentrations of 2 and 4 mg/ml collagen. For each condition, 50 µl of unpolymerized collagen solution was placed on glass-bottom dishes and incubated at 37°C and 5% CO_2_ for 1 hr. After the gels polymerized, 500 µl media were added to the wells.

### 2.3. Morphological assessment

Cellular morphology was assessed on Day 1 for isolated cells and on Days 4, 7 and 10 for cell clusters. Media was replaced every three days. Phase-contrast images of cells were acquired on a Zeiss LSM 880 with Airyscan confocal microscope (Carl Zeiss Microimaging, Germany) with a 63x 1.4 oil immersion objective and an electron-multiplying CCD camera (Hamamatsu Photonics, Hamamatsu, Japan). Cells or cell clusters located near the middle of the gel were imaged. At least 28 images were acquired per condition from three repeat experiments (n = 4).

Images were post-processed and morphometrically assessed using Image J [46] to quantify cell area A and roundness R = 4A/(π · L_M2_), where *L*_M_ refers to the length of the cell along the major axis. Higher values of *R* represented rounder shapes. Isolated cells and cell clusters were considered round for *R* ≥ 0.80 and elongated for *R* < 0.80.

### 2.4. Mitochondrial particle tracking microrheology

Microrheology experiments were conducted on isolated cells on Day 1 and cell clusters on Day 7. Mitochondria were fluorescently labelled 1 hr before experimentation by incubating cells with 350 nM Mitotracker Green solution (Life Technologies, Carlsbad, CA) added to RPMI 1640 media. Cells and cell clusters near the middle of the collagen gel were imaged in time-lapse mode (50 ms per frame for 120 s) to capture mitochondrial fluctuation. Time-lapse images were collected with a Zeiss LSM 880 with Airyscan confocal microscope (Carl Zeiss Microimaging, Germany) with 63x 1.4 oil immersion objective and an electron-multiplying CCD camera (Hamamatsu Photonics, Hamamatsu, Japan). Experiments were repeated at least once (n = 2), each assessing at least 24 isolated cells per condition and 16 cell clusters. At least four outer cells per cluster were assessed.

Time-lapse images of cells with at least 60 mitochondria visible for at least 80% of the total acquisition time were used for analysis. Mitochondrial particle trajectories were constructed in TrackMate [47] and converted to the mean square displacement (MSD) using @msdanalyzer [48] in MATLAB (The MathWorks, Natick, MA).

The ensemble-averaged MSD 〈Δr^2^ (τ)*_xy_*〉 was calculated according to:

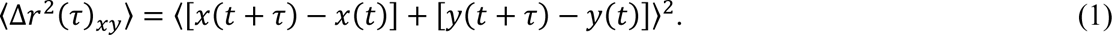

Here, τ is the delay time between the first and last image frame used for the analysis, and x(t) and y(t) are the spatial coordinates of mitochondrial particle positions at time t.

For viscoelastic materials, the MSD scales nonlinearly with τ according to a power-law relationship, 〈Δ*r*^2^ (τ)*_xy_*〉 ∼ τ^α^. The MSD-dependent power-law coefficient α was determined as a measure of intracellular fluidity (deformability):

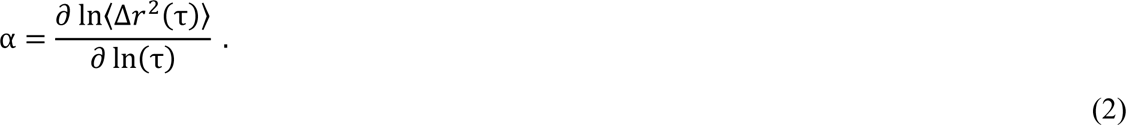

### 2.5. Immunofluorescent quantification of TBX3 and F-actin

Cells were stained for either TBX3 on Day 1 and 7 or F-actin on Day 7 as previously described [49]. In brief, cells were gently washed with PBS and fixed with 4% paraformaldehyde in PBS at room temperature for 20 min. Cells were washed thrice with PBS and permeabilized with 0.2% Triton X-100 in PBS at room temperature for 10 min. Cells were blocked in 5% sheep serum in PBS for 1 hr at room temperature and washed once with PBS/Tween. For TBX3 staining, cells on Day 1 and cell clusters on Day 7 were incubated overnight at 4°C with rabbit polyclonal anti-TBX3 antibody (Zymed 42-4800, Zymed Laboratories) at a 1:50 dilution in 5% sheep serum. Next, cells were washed thrice with PBS/Tween, incubated with Cy3 donkey anti-rabbit IgG (Jackson ImmunoResearch Laboratories, Inc., USA) at a dilution of 1:1000 for 60 min and washed thrice more with PBS/Tween. For actin staining, cell clusters on Day 7 were incubated overnight with 100 nM Acti-stain^TM^ 488 Phalloidin (Cat. #PHDG1, Cytoskeleton Inc., USA) and washed thrice with PBS. Nuclei were counterstained with 1 µg/ml Hoechst (Sigma, USA) for 10 min and washed thrice in PBS before imaging.

A series of confocal cross-sectional images with a constant step size of 0.5 µm (z-stack) per cell cluster was acquired (Zeiss LSM 880 AiryScan microscope, 63x 1.4 oil immersion objective) and converted to a single image employing the maximum intensity projection (MIP) (ZEN lite black edition, Zeiss, Germany). The fluorescence in MIP images was quantified as the corrected total cell fluorescence CTCF = *I* – *A*_ROI_ · *b*, where *I* is the integrated pixel density (sum of all pixels) of the region of interest for fluorescence quantification, *A*_ROI_ is the area of the region of interest, and *b* is the mean fluorescence of the background in the region of interest [50].

The schedule of experimental activities and assessments is summarized in supplemental Table S2.

### 2.6. Statistical analysis

The normality of data was evaluated using the Shapiro-Wilk test. Data were reported as mean ± standard deviation in the text and mean and standard error of the mean (SEM) in graphs. All evaluations used *p* < .05 for statistical significance unless otherwise stated, e.g., for multiple comparisons.

Regression models determined whether disease stage, collagen matrix concentration and time point of the experiments were predictors of the area and roundness of isolated cells and cell clusters. Multiple linear regression models were derived for area measurements to predict the cluster size as either absolute area *Â* or as area normalized to the mean single-cell area *Â*_N_ (normalized area). Beta regression models were employed to predict the roundness *Ȓ*.

Microrheology, TBX3 and F-actin data were assessed with multi-factorial ANOVA for significant main effects and two- or three-way interactions among variables. Significant interactions were further assessed with linearly independent pairwise comparisons among estimated marginal means.

Statistica for Windows (Version 13.5, TIBCO Software Inc., Palo Alto, CA) was used for regression model analysis, and SPSS for Windows (Version 25, IBM Corp., Armonk, NY) was used for all other analyses.

## 3. Results

### 3.1. Morphology

Morphology measurements were obtained from 28 images from three repeat experiments (n = 4).

#### 3.1.1. VGP isolated cells and cell clusters were larger than MET counterparts

For absolute cell area, the advanced disease stage (MET versus VGP) exhibited a marked decrease of the area *Â* on Day 1 and Day 4 (*p* < .001 and *p* = .001, respectively) and a slight decrease of *Â* on Day 7 and Day 10 (*p* = .254 and *p* = .168, respectively), Figure 1a. Increased collagen concentration from 2 to 4 mg/ml correlated with a decrease of *Â* of cells on Day 1 (*p* = .004). For cell clusters, increased collagen concentration correlated with a decrease of *Â* on Day 4 and 7 (*p* = .435 and *p* = .512, respectively), although this was not significant and had no effect on *Â* on Day 10 (*p* = .740).

**Figure 1:**
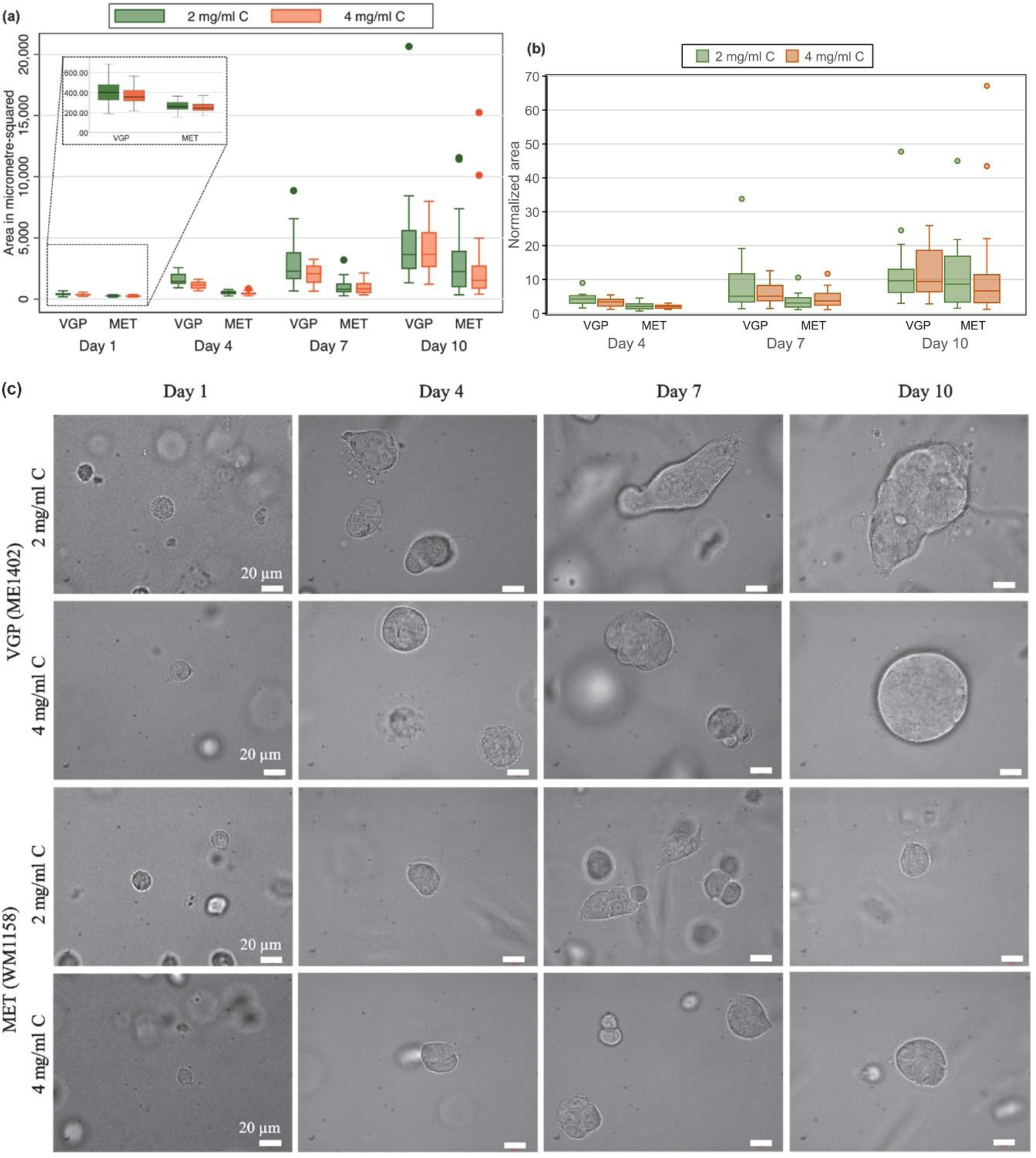
(a) VGP cells exhibited a larger area A than MET cells (p < .001) for all time points and collagen concentrations. (b) Normalized area of cell clusters for VGP was larger than for MET on Days 4 and 7 but similar on Day 10. Data for Day 1 as baseline for the normalization with Â_N_ = 1 is not shown. (c) Representative confocal brightfield images of VGP and MET cells in 2 and 4 mg/ml collagen concentration at Days 1, 4, 7 and 10. (All images 63x magnification and scale bars represent 20 µm.)

For normalized area, advanced disease stage correlated with a decrease in *Â*_N_ on Day 4 and Day 7 (*p* < .001 and *p* =.001, respectively) but did not alter *Â*_N_ on Day 10 (*p* = .222), Figure 1b. Increased collagen concentration correlated with a decrease in *Â*_N_ on Day 4 and Day 7 (*p* = .154 and *p* = .616, respectively), although not significant, and did not alter *Â*_N_ on Day 10 (*p* = .990).

Finally, increased time points from Day 1 to Day 4, 7 and 10, respectively, correlated with an increase of *Â* and *Â*_N_ (*p* < .001 for all cases), indicating cluster growth.

Pairwise comparisons indicated that the absolute area was larger for VGP cells than for MET cells at each time point and for both collagen concentrations (*p* < .001). However, when normalized, VGP clusters were larger than MET clusters on Days 4 and 7 in 2 mg/ml (*p* < .001 for both cases) and in 4 mg/ml collagen (*p* < .001 and *p* = .002, respectively). For Day 10, VGP and MET clusters had similar normalized area in 2 and 4 mg/ml collagen (*p* = .085 and *p* = .008, respectively). Bonferroni correction led to significance at *p* < .006.

#### 3.1.2. Roundness of cell clusters increased with increased collagen concentration

Isolated cells and cell clusters were considered round for median roundness *R* ≥ 0.80 and elongated for *R* < 0.80 (Figure 1b). Isolated VGP and MET cells (Day 1) were predominantly round, regardless of collagen concentration, with some notably elongated cells (outliers). In comparison, VGP and MET cell clusters were mostly elongated in 2 mg/ml collagen and round in 4 mg/ml collagen (Figure 2).

**Figure 2:**
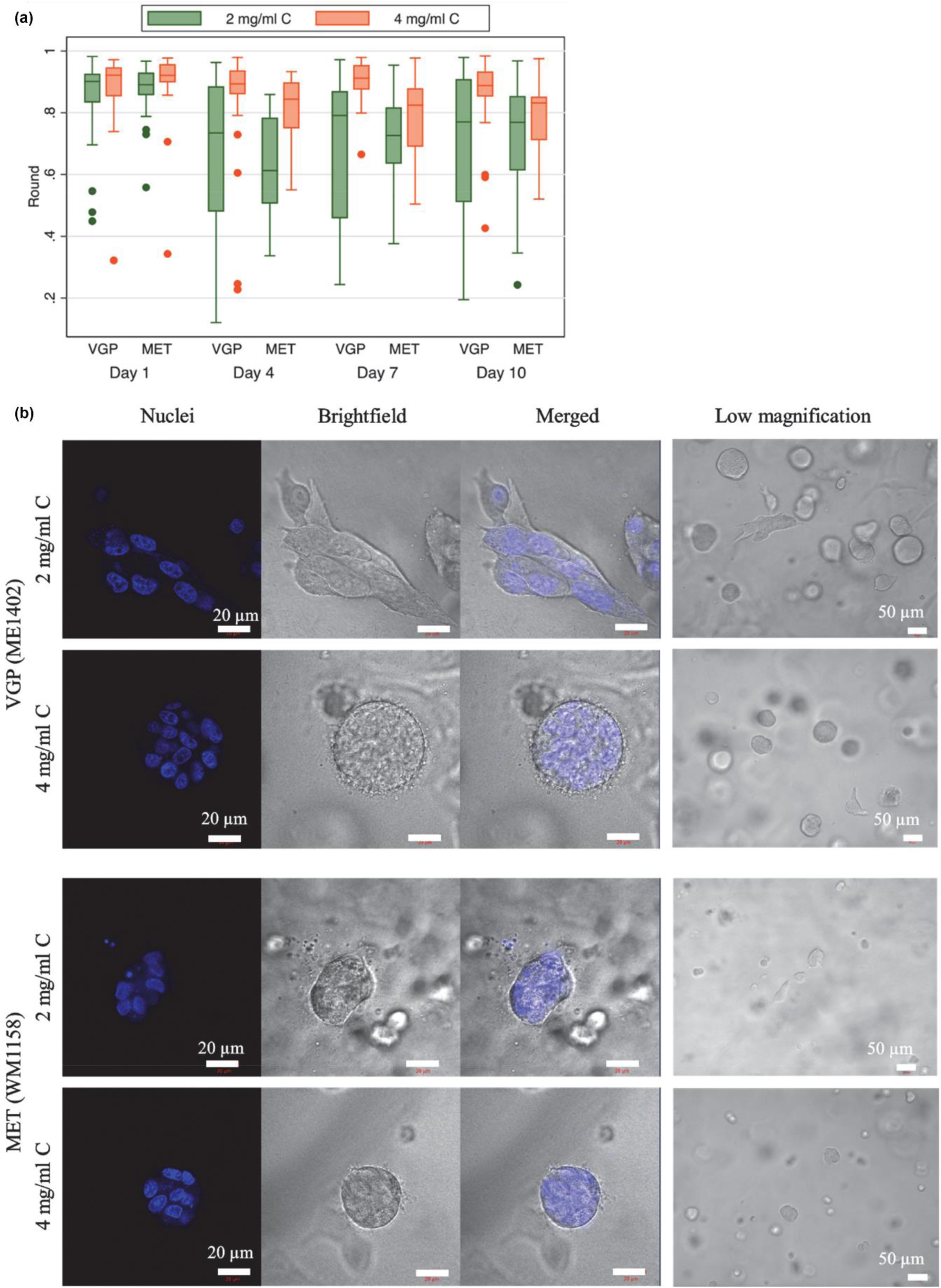
(a) Isolated VGP and MET cells on Day 1 were round (R ≥ 0.8) irrespective of the collagen concentration of the matrix. VGP and MET cell clusters were mostly elongated (R < 0.8) for 2 mg/ml collagen concentration (C) and round for 4 mg/ml collagen concentration. (b) Confocal images (2D confocal plane) of VGP and MET clusters after 7 days in 2 and 4 mg/ml collagen (C) matrices. Each panel presents (from left to right) fluorescently labelled nuclei (blue), a brightfield image, a merged image, and a low-magnification brightfield image showing multiple clusters. In 2 mg/ml C, the elongated shape was more defined for VGP clusters than MET clusters. Increased collagen concentration compacted and rounded the cell clusters. (Nuclei, brightfield and merged images 63x magnification, scale bars represent 20 µm. Low magnification images 20x magnification, scale bars represent 50 µm).

A beta regression model showed that an increase in time from Day 1 to 4, 7 and 10, respectively (*p* < .001, *p* = .002, and *p* = .006, respectively), correlated with a decrease in *Ȓ*. In contrast, an increase in collagen concentration from 2 to 4 mg/ml correlated with an increase in *Ȓ* on Day 4 and 7 (*p* =.045 and *p* = .019, respectively) but did not contribute to changes in *Ȓ* on Day 1 and Day 10 (*p* =.327 and *p* =.132, respectively).

### 3.2. Structural properties of cell clusters

The actin cytoskeleton is vital in controlling global cell morphology and structure [51]. Cell clusters on Day 7 were processed for immunofluorescence to detect F-actin and visualized by confocal microscopy.

#### 3.2.1. Elongated VGP clusters formed supracellular features, and large VGP clusters merged from smaller clusters

Elongated VGP clusters were categorized as sharp- or blunt-edged in 2 mg/ml collagen (Figure 3a, b). Sharp-edged clusters comprised mostly large, elongated cells that formed supracellular features, such as supracellular protrusive fronts and retracting tails (Figure 3a, blue arrows). Elongated clusters with a blunt edge showed no supracellular protrusions; however, microspikes were present. More importantly, 2D planes showed clear borders outlining at least two clusters that had merged into one cluster (see dashed lines in Figure 3b). Interestingly, actin visualization demonstrated merged VGP clusters in 2 mg/ml collagen (Figure 3c). Thick actin bundles (white arrow) were apparent at the interface of merged clusters.

**Figure 3:**
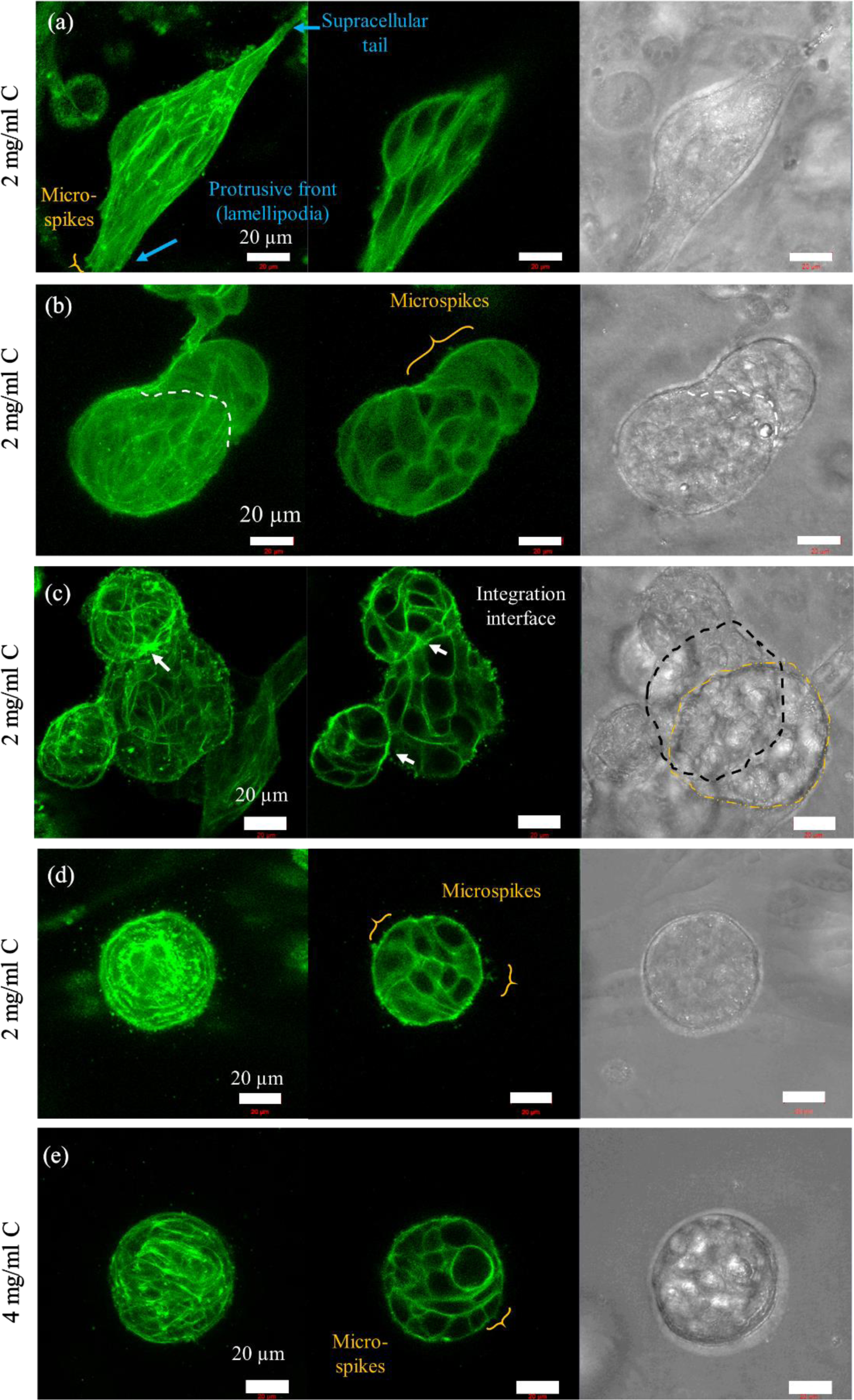
Confocal images of elongated (a, b), merged (c), and rounded (d, e) VGP cell clusters after 7 days in 2 and 4 mg/ml collagen (C) matrices. Each panel shows the mean intensity projection (MIP) of multiple fluorescently labelled F-actin (green) of confocal image stacks (left), F-actin of the mid-2D plane (middle) and MIP of brightfield confocal image stacks (right). All images at 63x magnification; scale bar represents 20 µm. (a,b) Sharp or blunt edges distinguished elongated VGP clusters in 2 mg/ml collagen: A sharp-edged cluster (a) exhibiting supracellular edge features, i.e. protrusive front and retracting tail (blue arrows and text). Microspikes (gold bracket) were visible on the protrusions. A blunt-edged cluster (b) merged from two smaller clusters (white dashed line shows the merging site). (c) Merged clusters resulted from two or more cell clusters that migrated towards each other and merged in 2 mg/ml collagen. The interface of merged clusters (white arrows) consisted of thick F-actin bundles in fluorescence images. The merged interface in (c) is outlined (black dashed curve) in brightfield images to distinguish it from the separate cluster not part of the merged cluster. The yellow dashed curve outlines a separate cluster that appeared in the field of view but is not attached to the merged cluster. (d, e) For round VGP clusters in 2 mg/ml collagen (d) and 4 mg/ml collagen (e), pseudopodia were primarily finger-like protrusions. Most round clusters in 4 mg/ml C contained a large cell towards the cluster interior.

Round VGP clusters exhibited similar actin-filled pseudopodia in 2 and 4 mg/ml collagen (Figure 3c, d). Several filopodia and microspikes emanated from cells at the cluster surface. Many clusters in 4 mg/ml collagen comprised a single large and round cell (R > 0.90) with a thick actin cortex in the cluster’s interior. This cell was more than twice the size of neighbouring cells.

#### 3.2.2. MET clusters exhibited several types of pseudopodia in 2 mg/ml collagen but were only in the form of filopodia in 4 mg/ml collagen

MET clusters invaded the 2 mg/ml collagen environment as either loose (i.e. non-adherent) or compact cell groups. In loose MET clusters, the first cell in the cluster, the leader cell, had a planar protrusive front. All cells had a retracting tail (see blue arrows in Figure 4a). Each cell within the cluster exhibited a thick actin cortex and filopodia and microspikes emanating from the cell membrane. The thickest actin cortex and the most prominent filopodia and microspikes were observed in the leader cell (Figure 4a).

**Figure 4:**
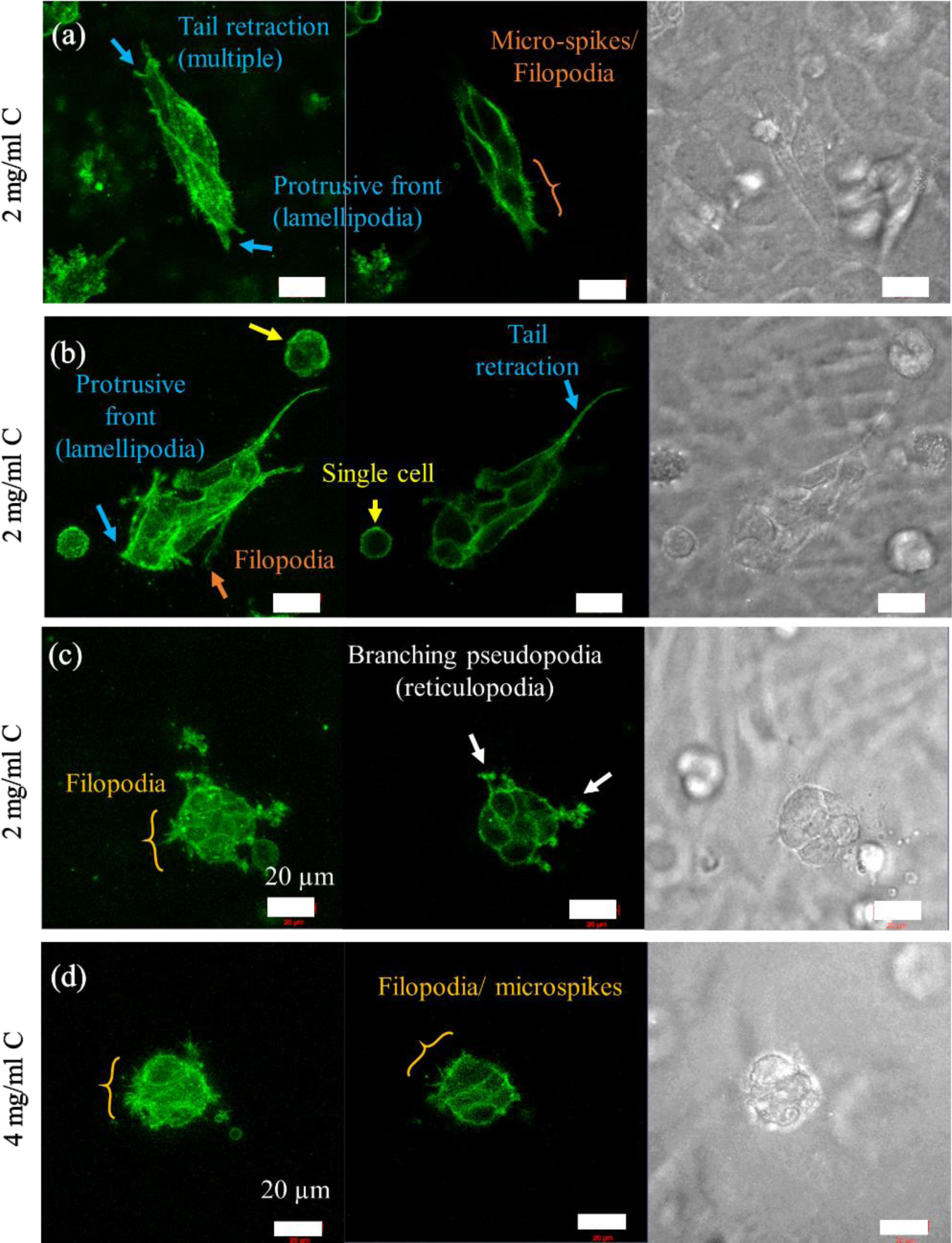
Confocal images of elongated (a, b) and round (c, d) MET cell clusters after 7 days in 2 and 4 mg/ml collagen (C) matrices. Each panel shows the mean intensity projection (MIP) of multiple fluorescently labelled F-actin (green) of confocal image stacks (left), F-actin of the mid-2D plane (middle) and MIP of brightfield confocal image stacks (right). All images at 63x magnification; scale bar represents 20 µm. (a, b) Elongated MET clusters were loose or compact in 2 mg/ml collagen. (a) Loose clusters comprised cells aligned with each other. Each cell in the cluster featured one retracting tail (blue arrow and label). (b) Compact elongated clusters emerged as supracellular units with a protrusive front and retracting tails (blue arrows and label). Single cells in the cluster’s vicinity were polarised, as indicated by the focalized F-actin in the cortex (yellow arrow and label). Slender filopodia and microspikes are indicated with orange arrows, respectively. (c, d) Rounded MET clusters exhibited complex branching pseudopodia (white arrows) and filopodia (orange bracket) in 2 mg/ml collagen (c) compared to simple filopodia and microspikes in 4 mg/ml collagen (d). Related quantitative data are shown in Figure 2a.

Compact MET clusters were either elongated or round in 2 mg/ml collagen, as in VGP clusters, and primarily round in 4 mg/ml collagen. Elongated compact clusters formed supracellular planar protrusive front and retracting tails (Figure 4b). Round MET clusters extended varying pseudopodia with a complexity that depended on matrix rigidity. In 2 mg/ml collagen, round MET clusters exhibited branching filamentous pseudopodal networks, known as reticulopodia, and several filopodia (Figure 4c). In 4 mg/ml collagen, long filopodia and short microspikes were present (Figure 4d).

### 3.3. Mechanical properties

Mitochondrial tracking experiments were conducted to investigate the intracellular mechanics of isolated cells and cell clusters. Mitochondrial fluctuation was traced and used to calculate the mean squared displacement (MSD) and power-law coefficient α.

Low MSD values, based on small particle fluctuation, correspond to inhibited mitochondrial displacement and indicate stiffer, solid materials. High MSD values, representing large mitochondrial fluctuation, indicate greater particle motility and a softer, more fluid-like material. The MSD was determined for different delay times τ, i.e. the period between the first and the last image frame used for the analysis. Particles may displace either small lengths due to random forces from the surrounding medium’s thermal energy during short delay times or large lengths due to active, motor-driven processes at long delay times [2, 44, 52].

The power-law coefficient α (representing the slope of the logarithmic MSD-delay time curve) characterizes the power-law delay time dependence and helps classify the tracer particles’ motion. A value of α close to 1 corresponds to diffusive motion, such as thermal fluctuation in a Newtonian fluid, whereas a value of α close to 0 indicates constrained, sub-diffusive motion, such as thermal fluctuation in an elastic material [44, 53]. Mitochondrial particles typically experience considerable intracellular traffic and therefore undergo constrained, sub-diffusive particle fluctuation with 0 < α < 1, which can be further classified into strong (0 < α < 0.3) or weak (0.7 < α < 1) sub-diffusive motion.

Lower and flat MSD-τ curves (e.g. Figure 5 b-d) indicate solid-like intracellular behaviour, whereas higher and increasingly sloped curves indicate more fluid-like, intracellular motor-driven mechanics.

**Figure 5:**
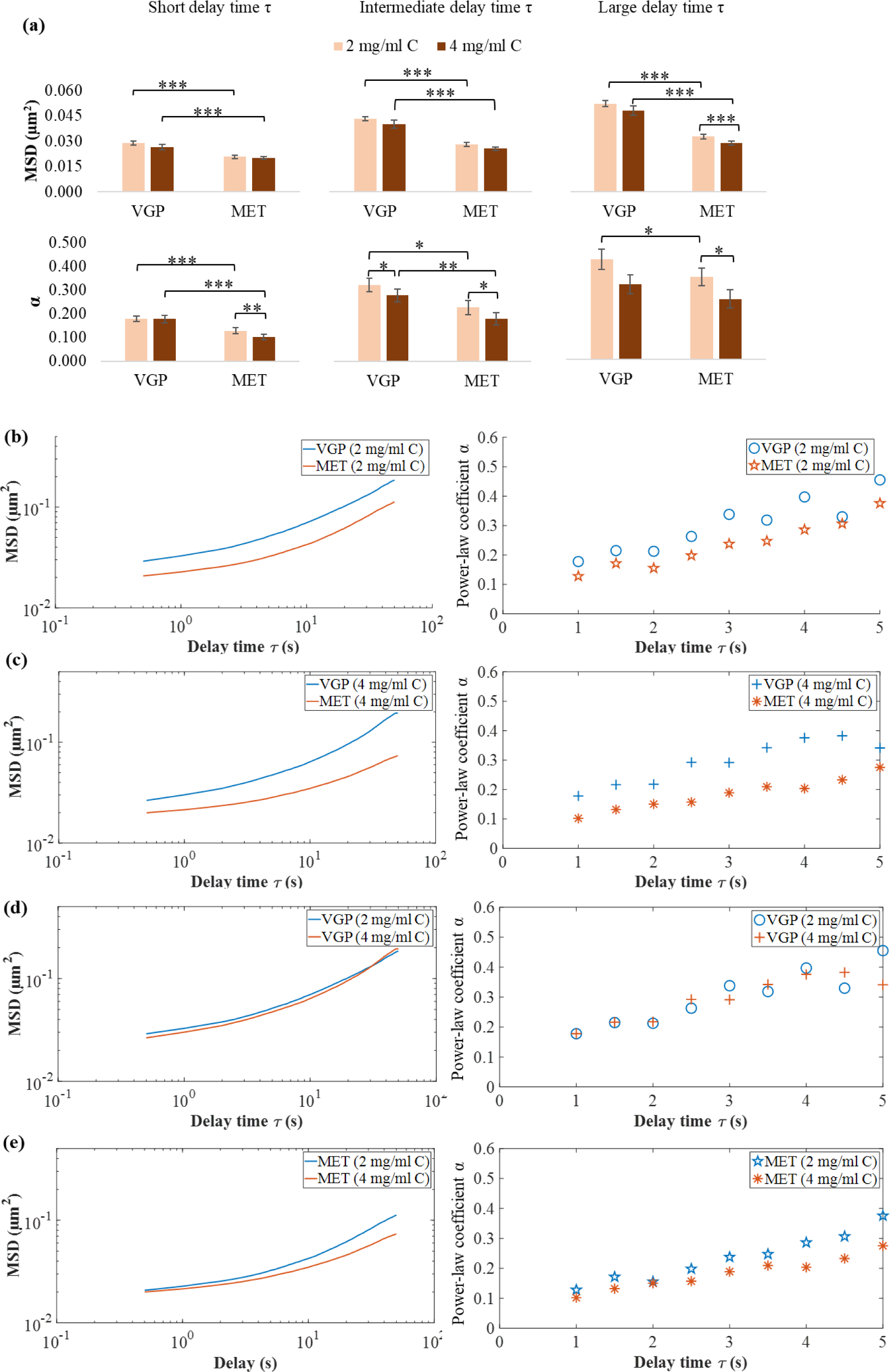
MSD and power-law coefficient α of VGP and MET **isolated cells** (Day 1) in matrices with 2 and 4 mg/ml collagen concentration (C). a) Comparison of disease stage and collagen concentration at short (MSD: τ = 0.05 s, α: τ = 1 s), intermediate (MSD and α: τ = 3 s) and long delay time (MSD and α: τ = 5 s). b, c) Comparison of different disease stages for the same collagen concentration. MET cells had lower MSD and α than VGP cells in 2 mg/ml C (b) and 4 mg/ml C (c). d, e) Comparison of the same disease stage for different collagen concentrations. MSD and α of VGP cells were similar in 2 and 4 mg/ml C (d), whereas MSD and α of MET cells were higher in 2 than 4 mg/ml C (e). Higher MSD and α indicate increased mitochondrial fluctuation and decreased cell stiffness. Error bars were omitted for clarity.

#### 3.3.1. Isolated cells were more deformable at VGP than MET stage

For isolated cells, the MSD and power-law coefficient α were larger for VGP cells than for MET cells in 2 and 4 mg/ml collagen matrices at short (MSD at τ = 0.05 s and α at τ = 1 s), intermediate (MSD and α at τ = 3 s), and long delay times (MSD and α at τ = 5 s), see Figure 5(a) and Table S1.

The MSD and α of isolated VGP cells were similar in 2 and 4 mg/ml collagen matrices for all delay times τ. For MET cells, the MSD of cells in 4 mg/ml collagen compared to cells in 2 mg/ml collagen increased with delay times, with statistical significance indicated for τ ≥ 5 s. α of MET cells decreased for increased collagen concentration for all delay times (Figure 5a).

A decrease in MSD and α indicated that mitochondrial fluctuation was impeded and cells changed from fluid- to solid-like behaviour, respectively. The more advanced MET disease stage exhibited decreased mitochondrial fluctuation (i.e. MSD) and increased cell stiffness of isolated cells in 2 and 4 mg/ml collagen (Figure 5b - e). Increased collagen concentration correlated with decreased mitochondrial fluctuation and increased cell stiffness for MET but not VGP cells.

#### 3.3.2. VGP and MET cell clusters exhibited similar deformability

The MSD of cell clusters was larger at the MET than the VGP disease stage in 2 mg/ml collagen for short, intermediate, and large delay times. However, the α of VGP and MET clusters were similar. In 4 mg/ml collagen, the MSD and α of VGP and MET clusters were similar.

For VGP clusters, the MSD increased for all τ, whereas α decreased with increased collagen concentration. A significant reduction in α was detected for τ ≥ 3s. A reduction in α indicated that increased collagen concentration stiffened or induced a solid-like response in VGP cells. For the MET stage, the MSD and α of clusters in 2 and 4 mg/ml collagen were similar (Figure 6a).

**Figure 6:**
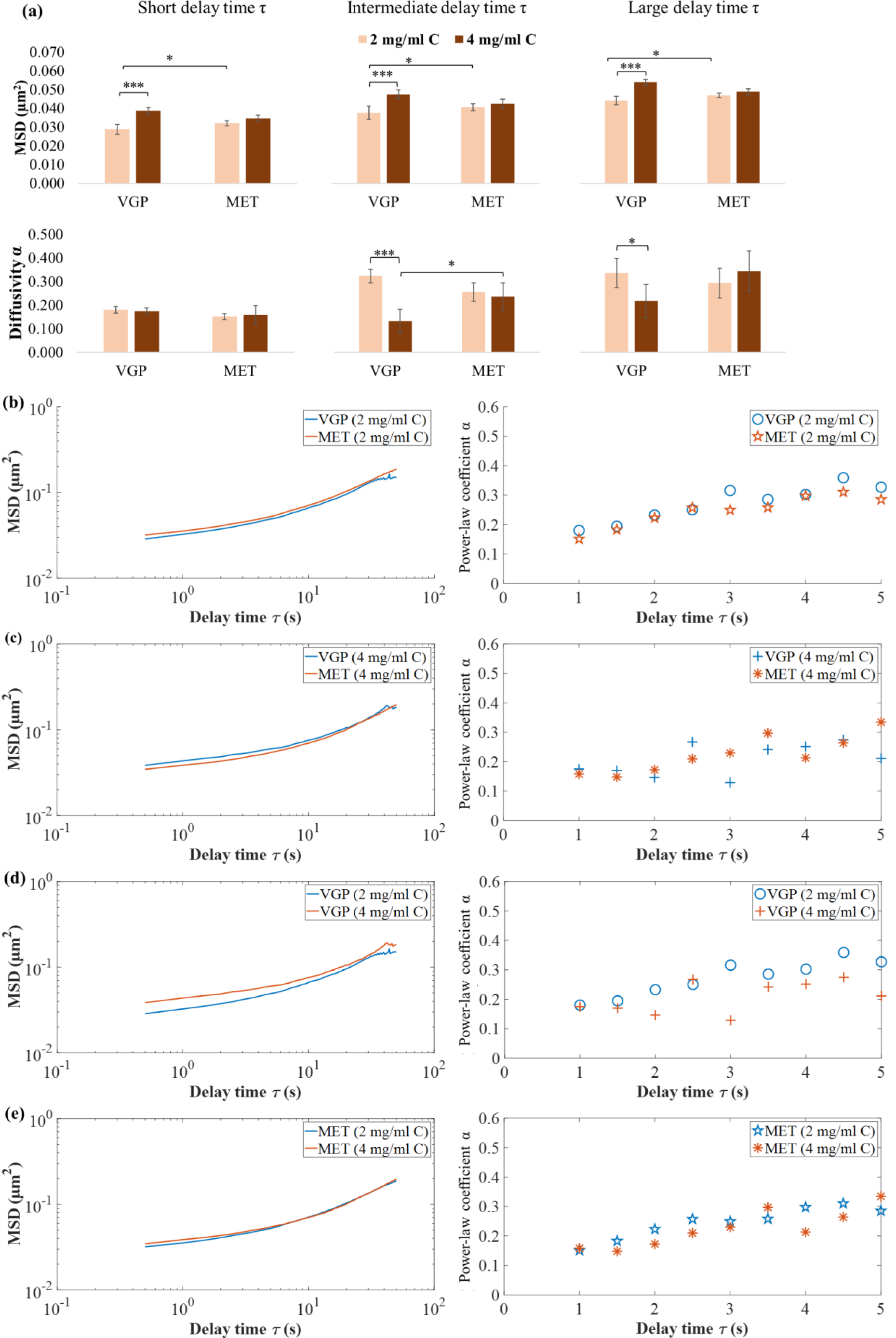
MSD and power-law coefficient α of VGP and MET **cell clusters** (Day 7) in matrices with 2 and 4 mg/ml collagen concentration (C). a) disease stage and collagen concentration at short (MSD: τ = 0.05 s, α: τ = 1 s), intermediate (MSD and α: τ = 3 s) and long delay time (MSD and α: τ = 5 s). b, c) Comparison of different disease stages for the same collagen concentration. The MSD and power-law coefficient α of VGP clusters were similar to that of MET clusters in 2 mg/ml C (b) and 4 mg/ml C (c). d, e) Comparison of the same disease stage for different collagen concentrations. For VGP clusters, the MSD was smaller, and α was larger in 2 mg/ml C than in 4 mg/ml C (d). MET clusters did not exhibit differences in MSD and α in 2 and 4 mg/ml C (e).

The advanced disease stage of cell clusters was reflected in increased mitochondrial fluctuation (MSD) in 2 mg/ml collagen but did not change in 4 mg/ml collagen. The cell stiffness of MET and VGP clusters was similar both in 2 mg/ml and 4 mg/ml collagen (Figure 6b - e). Increased collagen concentration increased mitochondrial fluctuation and cell stiffness in VGP clusters but not in MET clusters.

#### 3.3.3. Mitochondrial fluctuation increased with cluster formation, except for VGP cells in soft matrices

Active processes such as molecular motor activity, prevalent during larger delay times [2], typically drive the mitochondrial fluctuation into diffusive regimes [52]. Our results did not indicate significant effects of such active processes for the selected delay times since the mitochondrial fluctuation of cells before and after cluster formation were strongly sub-diffusive (α ≤ 0.3).

Comparison of MSD data during the onset of cluster formation in 2 mg/ml collagen revealed that the MSD and α of VGP clusters (Day 7) were similar to that of isolated VGP cells (Day 1) for all τ (Figure 7a). For MET cells, the MSD increased, but α did not change while isolated cells evolved into cell clusters (Figure 7b).

**Figure 7:**
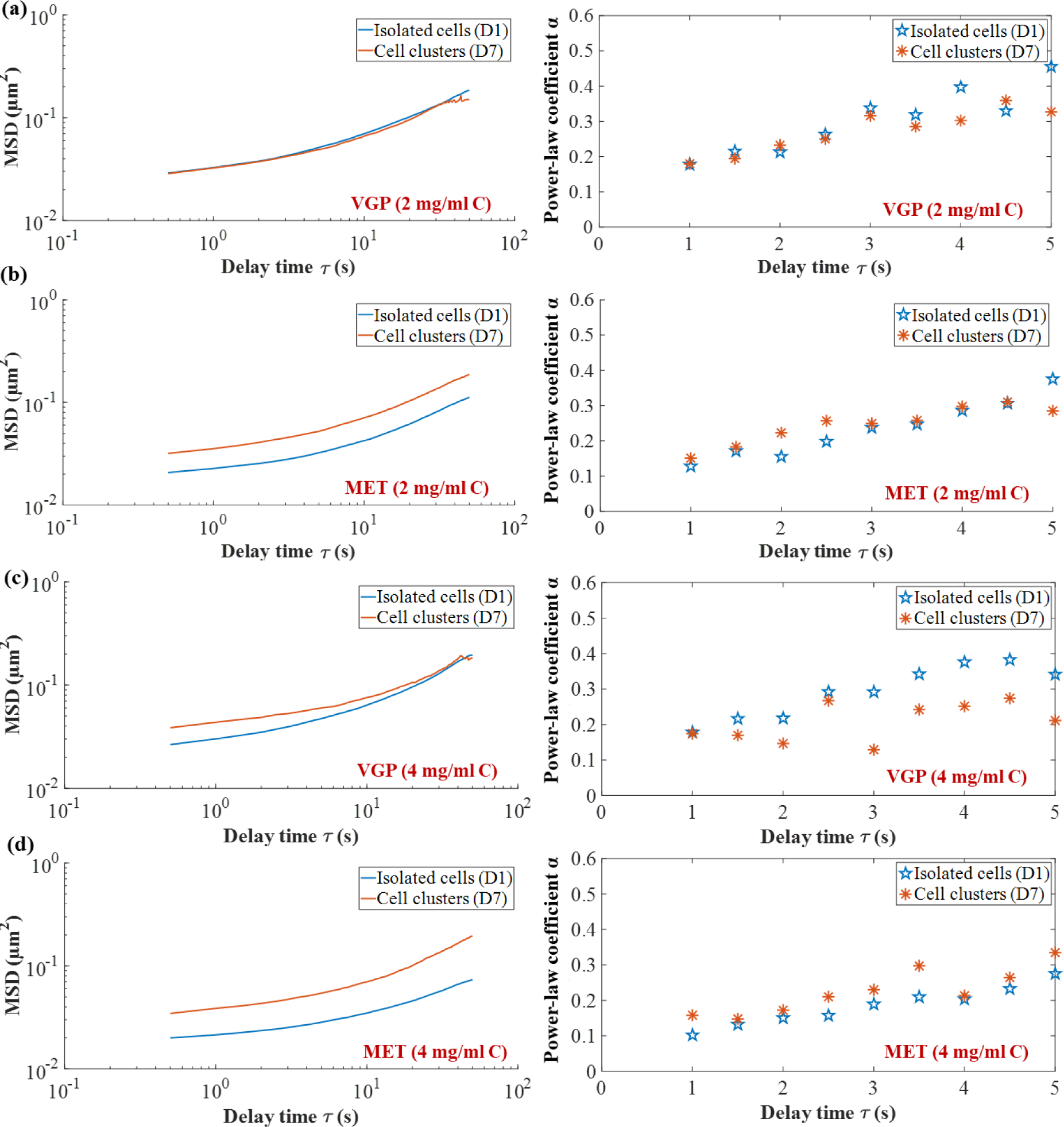
MSD and power-law coefficient α of **isolated cells** and **cell clusters** in 2 mg/ml (a, b) and 4 mg/ml collagen (C) matrices (c, d) to illustrate the effect of cluster formation on intracellular microrheological properties. In 2 mg/ml collagen, the MSD and α of VGP isolated cells were similar to those of VGP cell clusters (a). For MET cells, the MSD was larger for clusters than for isolated cells, whereas α did not change (b). In 4 mg/ml collagen, VGP clusters exhibited larger MSD and α than isolated VGP cells (c). For MET cells, MSD substantially increased, but α decreased during cluster formation (d). (D1 = Day 1 and D7 = Day 7. Error bars omitted for clarity.)

In 4 mg/ml collagen, the MSD was larger for VGP clusters than for isolated VGP cells for τ < 10 s. However, the MSD converged for the longest delay times considered. The power-law coefficient α of VGP clusters was smaller than that of isolated VGP cells for τ > 1 s (Figure 7c). The MSD of MET cells increased with cluster formation, but unlike VGP cells, diverged for the longest delay times. The power-law coefficient α of MET clusters was larger than that of isolated MET cells for all τ (Figure 7d). Decreased α in VGP and increased α in MET cells indicated that VGP cells stiffened, whereas MET cells softened during cluster formation.

In summary, the onset of cluster formation in 2 mg/ml collagen led to increased mitochondrial fluctuation of MET cells but did not affect VGP cells. Further, similar α indicated that cells were equally stiff before and after cluster formation. In contrast, cluster formation in 4 mg/ml collagen generally led to increased mitochondrial fluctuation of VGP and MET cells. However, the cell stiffness of VGP cells increased, whereas that of MET cells decreased (Figure 7).

### 3.4. Expression of oncogenic factor TBX3

TBX3 belongs to the T-box family of transcription factors with fundamental roles in development, cell fate and tissue homeostasis [54]. However, overexpression of TBX3 has been implicated in several cancers where it is crucial for cell migration and tumour formation [43].

For immunofluorescence assessment, TBX3 of at least nine isolated cells (Day 1) and nine cell clusters (Day 7), for n = 2 experiments, were stained, and cell nuclei were counterstained (Figure 8). Nuclear TBX3 was measured.

**Figure 8:**
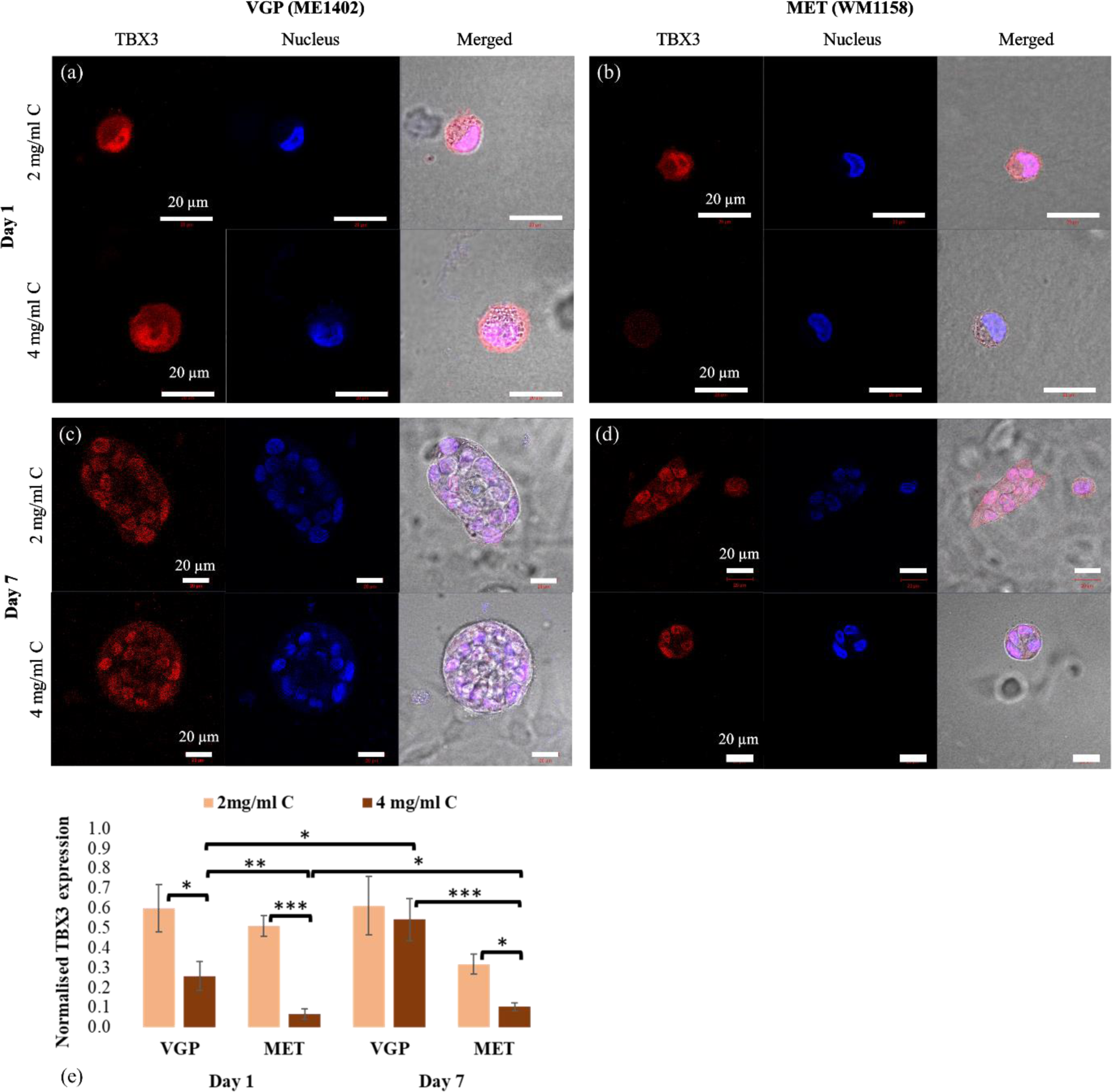
Fluorescent confocal images of TBX3 (red, left column), nucleus (blue, middle column) and merged TBX3 and nucleus superimposed on brightfield images (right column) for VGP and MET cells in 2 and 4 mg/ml collagen matrices after 24 hrs and 7 days (a – d). In merged TBX3 and nucleus fluorescence, pink indicates colocalization of TBX3 in the nucleus. (a, b): For isolated cells, TBX3 was expressed in the nuclei of VGP cells in 2 and 4 mg/ml collagen (a) and the nuclei of MET cells in 2 mg/ml collagen but not in 4 mg/ml collagen (b). (c, d): For cell clusters, TBX3 was expressed in the nuclei of VGP clusters in 2 and 4 mg/ml collagen (c) and nuclei of MET clusters in 2 and 4 mg/ml collagen (d). For each condition, the sample size was N = 9 from n = 2 repeat experiments. Mid-2D focal plane, 63x magnification, scale bar represents 20 µm. *(e): Fluorescence of TBX3 normalized to nuclei. For isolated cells (Day 1), TBX3 was highly expressed in VGP and MET cells in 2 mg/ml collagen but decreased with increased collagen concentration. In 4 mg/ml collagen, TBX3 was expressed in isolated VGP cells but absent in MET cells. For cell clusters (Day 7), TBX3 in VGP was equally highly expressed in 2 and 4 mg/ml collagen. In MET clusters, TBX3 was expressed but lower in 4 mg/ml than 2 mg/ml collagen. (Statistical analyses were performed on the logarithmic transformation of normalized TBX3 data since untransformed data were not normally distributed; *** p < .001, ** p < .005 and * p < .05.)*

#### 3.4.1. TBX3 in isolated cells was highly expressed in soft but not in stiff matrices

TBX3 was equally highly expressed in isolated VGP and MET cells in 2 mg/ml collagen (*p* = .987) (Figure 8e, Day 1). Compared to 2 mg/ml collagen, in 4 mg/ml collagen, cellular TBX3 levels decreased by 57% (*p* = .007) for VGP and by 87% (*p* < .001) for MET cells. Notably, TBX3 was still detectable and present in VGP cells but nearly disappeared in MET cells in 4 mg/ml collagen.

#### 3.4.2. TBX3 decreased with increased collagen concentration in MET but not VGP cell clusters

In cell clusters, TBX3 levels for MET were 48% lower than for VGP in 2 mg/ml collagen (*p* = .344, non-significant) and 81% lower than for VGP in 4 mg/ml collagen (*p* < .001) (Figure 8e, Day 7). TBX3 of MET clusters was 68% lower in 4 mg/ml than in 2 mg/ml collagen (*p* = .006), whereas TBX3 in VGP clusters was equally highly expressed in 2 and 4 mg/ml collagen (*p* = .877).

#### 3.4.3. TBX3 increased during cluster formation in stiff but not soft matrices

TBX3 levels during VGP cluster formation (i.e. transition from isolated cells at Day 1 to cell clusters at Day 7) increased for 4 mg/ml collagen (*p* =.028) but did not change for 2 mg/ml collagen (*p* =.731) (Figure 8e). For MET cluster formation, TBX3 levels increased in 4 mg/ml collagen (*p* = .032) and decreased in 2 mg/ml collagen (*p* = .201, non-significant).

## 4. Discussion

We quantified and compared *in vitro* the morphology, intracellular mechanics and TBX3 expression in VGP and MET melanoma cells in isolation and cell aggregates following cluster formation over 10 days in 3D collagen matrices of two different rigidities.

It was found that the intracellular stiffness of VGP and MET cells changed during cluster formation in stiff but not soft matrices, indicating that soft matrices provide a more favourable environment for tumour growth. High TBX3 expression was observed in isolated VGP and MET cells in soft matrices. However, VGP cells rapidly formed cell clusters that increased in size over time, whereas MET cells remained predominantly isolated. This difference in cluster formation suggests that high TBX3 levels mediate cluster formation and collective cell migration modes at the VGP disease stage but play a lesser role in the more advanced metastatic melanoma stage.

### 4.1. Isolated cells

Compared to VGP cells, MET cells exhibited smaller volume and mitochondrial fluctuation (i.e. MSD) but higher stiffness in soft and stiff 3D collagen matrices. This reflects evolutionary changes associated with melanoma disease progression. Regulation of cell volume has important biomechanical and biochemical implications. For example, reports have shown that reduced cell volume increases cell stiffness [55] and molecular crowding [56], hindering protein mobility and slowing several biochemical cascades. Importantly, we observed increased stiffness of melanoma cells in the more advanced MET disease stage than in the VGP stage. This contrasts several studies that report decreased stiffness with melanoma disease progression [57-59]. However, these studies did not consider the effect of the change in cell volume. Few studies have reported increased stiffness with increased metastatic potential in prostate [60, 61] and melanoma cancers [62].

It is important to note that VGP cells formed clusters, whereas MET cells predominantly invaded the 3D collagen matrices in isolation. It is recognized that the size and deformability of cancer cells may facilitate penetration through several barriers for secondary colonization [57]. However, VGP cells are yet to penetrate thick layers of skin and need mechanical reinforcement to do so [63]. It is proposed that the deformability of the large VGP cells is insufficient to allow isolated cells to invade the dermis efficiently. Instead, large cluster formation may serve as a more efficient alternative to invade the surrounding tissue than single-cell invasion.

Our results showed high TBX3 expression in isolated VGP and MET cells in soft matrices. As previously mentioned, VGP cells quickly formed cell clusters, whereas MET cells mostly remained as isolated cells over time. Considering that TBX3 promotes tumour formation and metastasis [43], it is suggested that highly expressed TBX3 promotes migration and cluster formation in isolated VGP cells and migration but not cluster formation in MET cells.

A polyclonal antibody was used for TBX3 staining since TBX3 has previously been identified in the cytoplasm of cancer cells and it is thought that in addition to functioning as a transcription factor in the nucleus, it may have additional roles in the cytoplasm. A monoclonal antibody may have provided better staining if it is assumed that the cytoplasmic staining is non-specific.

Compared to the highly expressed intracellular TBX3 in soft matrices, TBX3 content was 2.3-fold lower in VGP cells and absent in MET cells in the stiff collagen matrix. These results indicate that TBX3 is partly governed by the cell’s mechanical interaction with its environment and the crucial signalling molecules [43]. Specifically, it is proposed that increased matrix rigidity increases actomyosin contractility, which stimulates signalling molecules that have overlapping functions in cytoskeletal remodelling and the PI3K/AKT signalling pathway. Actomyosin contractility is primarily mediated by Rho GTPases, such as RhoA, Rac1 and Cdc42 [64]. Interestingly, it has been reported that Cdc42 activity decreases with increased collagen matrix rigidity [65] and that upregulation of the PI3K/AKT pathway correlates with decreased Cdc42 levels [66].

Cell stiffness increased for isolated MET cells but did not change for VGP cells with increased matrix rigidity. Recent studies have shown that increased matrix rigidity through increased collagen concentration contributes to decreased cell stiffness of isolated prostate [67] and breast cancer cells [44] in 3D environments. However, using cell lines of breast, colon, and pancreas cancers of various stages, Wullkopf et al. [18] showed that the cell stiffness of early-stage cancer cells was consistently unaffected by changes in matrix rigidity. In contrast, the cell stiffness of highly metastatic cells either increased or decreased, even within the same cancer type. For example, the cell stiffness of metastatic breast cancer MDA-MB-231 cells increased, whereas the stiffness of 4T1 metastatic breast cancer cells decreased with increased matrix rigidity [18].

The VGP cells’ inability and the MET cells’ ability to adjust their intracellular biomechanics in response to external cues agree in part with the results of Wullkopf et al. [18]. However, it is essential to note that the dissimilarity of VGP and MET cells in adjusting their mechanical properties may be rooted in the functional interaction between cell volume and the extracellular environment. In soft matrices, VGP cells were approximately double the size of MET cells. Accordingly, increased matrix rigidity condensed the volume of large VGP and small MET cells by 11.4 and 5.9%, respectively. Due to the large size, the VGP cells’ compression may not immediately affect intracellular traffic, mitochondrial fluctuation, and cell stiffness. In contrast, intracellular traffic was innately confined in small MET cells in soft matrices. Condensation of MET cell size through increased matrix rigidity further compromised intracellular traffic.

### 4.2. Cluster formation and cell clusters

The onset of cluster formation for VGP cells was associated with changes in intracellular mechanical properties in rigid matrices but not in compliant matrices. In the soft collagen matrices, the mitochondrial fluctuation and cell stiffness of VGP cell clusters (Day 7) were the same as that of isolated cells (Day 1). This steadiness of the intracellular properties suggests that VGP cells were innately endowed with suitable mechanical properties to form clusters in soft matrices. Cluster formation appeared as an automatic cell fate of VGP cells.

The similarity of TBX3 levels in isolated VGP cells and VGP cell clusters supports the innate endowment of VGP cells with essential internal machinery for cluster formation. Based on the established role of TBX3 in promoting tumour formation and metastasis [43], we propose that high TBX3 expression in VGP cell clusters was imperative to mediate their collective migratory modes and the merging with adjacent clusters (see Figure 3). Further, VGP clusters in soft and stiff matrices were similar in cluster size (although different in shape) and, more importantly, in TBX3 expression. Remarkably, Li et al. [68] demonstrated a strong correlation between TBX3 expression and tumour size.

Our findings of increased mitochondrial fluctuation and cell stiffness during VGP cluster formation, i.e. compared to isolated cells, are substantiated by previous studies. Mih et al. [69] showed that increased matrix rigidity fosters increased proliferative conditions for the cell, and Cheng et al. [70] associated increased proliferation with increased mitochondrial activity.

In stiff matrices, VGP and MET cell clusters exhibited increased TBX3 expression compared to low and absent TBX3 expression, respectively, in isolated VGP and MET cells. Thus, increased expression demonstrates that TBX3 is indispensable during cluster formation in stiff environments.

It is also suggested that TBX3 enhances cluster formation through cell proliferation in stiff environments, considering that these environments facilitate an increased proliferative activity of cells [69]. Previous studies have argued that TBX3 promotes cell migration but impedes proliferation in pancreatic ductal adenocarcinomas, melanoma and breast cancers in 2D *in vitro* and in *in vivo* animal environments [71]. However, 2D *in vitro* conditions have limited physiological relevance [44]. Cells exhibit excessive proliferation and cell spreading in 2D compared to 3D environments, stemming from upregulated biochemical signalling that coordinates cell cycling and metabolism [4]. Animal models better represent human physiological conditions [44] but pose challenges for the controlled variation of matrix rigidity and differentiation of metastatic dissemination mode [45].

Advanced melanoma metastasizes to several organs in the body, including the lung, liver, brain and bone, with different environmental stiffnesses. Our finding that the intracellular stiffness changed during cluster formation in stiff but not soft matrices suggests that soft matrices provide a more favourable environment for cluster growth. Interestingly, this aligns with clinical and biopsy reports indicating that lungs with soft tissue are often the first and the most common distant melanoma metastasis target [72]. Differences in TBX3 expression in melanoma cells in 3D environments of different stiffness observed in this study may have important implications for developing therapeutic approaches. Melanoma therapies that inhibit high TBX3 expression may be effective in soft lung tissue but prove ineffective in stiff bone tissue. The change of intracellular stiffness and TBX3 expression in melanoma cells during cluster formation in differently stiff environments demonstrate the need to consider mechanical and molecular biomarkers of cancer malignancy in combination in a tissue-specific context.

In the current study, collagen concentration was used as a proxy for the stiffness of the collagen matrix. This approach has limitations, and it will be beneficial in future studies to quantify the matrix stiffness associated with different collagen concentrations to determine the alignment with physiological conditions. Collagen concentration also affects the matrix’s microarchitecture, which can vastly alter cell behaviour independent of matrix stiffness. Therefore, future approaches should incorporate the impact of collagen architecture and the degradation of the collagen matrix affecting the microarchitecture over time on cell behaviour. Future studies should also investigate the expression of TBX3 under different biomechanical conditions.

## 5. Conclusions

Advanced melanoma metastasises to several organs in the body, including lung, liver, brain and bone, each having a unique environmental stiffness. Our finding that the intracellular stiffness changed during VGP and MET cluster formation in stiff but not in soft matrices suggests that soft matrices provide a more favourable environment for tumour growth. Interestingly, this aligns with clinical and biopsy reports indicating that lungs with soft tissue are often the first and the most common distant melanoma metastasis target [72]. Differences in TBX3 expression in melanoma cells in 3D environments of different stiffness observed in this study may have important implications for developing therapeutic approaches. Melanoma therapies that inhibit high TBX3 expression may be effective in soft lung tissue but prove ineffective in stiff bone tissue. The change of intracellular stiffness and TBX3 expression in melanoma cells during tumour formation in differently stiff environments demonstrate the need to consider mechanical and molecular biomarkers of cancer malignancy in combination in a tissue-specific context.

## Data

Data supporting the results presented in this article are available on the University of Cape Town’s institutional data repository (ZivaHub) under http://doi.org/10.25375/uct.14254394 as Higgins G, Higgins F, Peres J, M. Lang D, Abdalrahman T, H. Zaman M, Prince S, Franz T. Data for “Intracellular stiffness, mitochondrial activity and TBX3 expression jointly dictate the spreading mode of melanoma cells in 3D environments”. ZivaHub, 2022, DOI: 10.25375/uct.14254394

## Supporting information

Supplement Table S1 Results comparisons

Supplement Table S2 Experimental design

## Acknowledgements

We acknowledge the technical assistance of Mrs Susan Cooper of the Confocal and Light Microscope Imaging Facility of the University of Cape Town. The microscope used in this work was purchased with funds from the Wellcome Trust (grant number 108473/Z/15/Z) and the National Research Foundation of South Africa (grant number UID93197). Ms Anneli Hardy from the Statistical Consulting Service at the Department of Statistical Sciences, University of Cape Town, is acknowledged for conducting the regression model analysis.

