## Supplement Table S1 Results comparisons for "Intracellular mechanics and TBX3 expression jointly dictate the spreading mode of melanoma cells in 3D environments"

Table S1: Adjustment of intracellular properties of VGP and MET melanoma cells in 2 and 4 mg/ml collagen (C) matrices. Mitochondrial fluctuations, intracellular stiffness, and TBX3 levels are represented by the mean square displacement (MSD) of mitochondria, the inverse of the diffusivity coefficient (α) and the signal intensity of TBX3 protein in the nucleus, respectively. Day 1 and Day 7 represent isolated cells and cell clusters, respectively.

| State of cell | Comparison | Conditions | MSD | α | TBX3 |
| --- | --- | --- | --- | --- | --- |
| Isolated cells only | VGP vs MET | 2 mg/ml C, Day 1 | Decrease | Decrease | Similar |
|  | VGP vs MET | 4 mg/ml C, Day 1 | Decrease | Decrease | Decrease |
| Cell clusters only | VGP vs MET | 2 mg/ml C, Day 7 | Similar | Similar | Decrease |
|  | VGP vs MET | 4 mg/ml C, Day 7 | Similar | Similar | Decrease |
| Isolated cells only | 2 vs 4 mg/ml C | VGP, Day 1 | Similar | Similar | Decrease |
|  | 2 vs 4 mg/ml C | MET, Day 1 | Decrease | Decrease | Decrease |
| Cell clusters only | 2 vs 4 mg/ml C | VGP, Day 7 | Similar | Similar | Similar |
|  | 2 vs 4 mg/ml C | MET, Day 7 | Similar | Similar | Decrease |
| Cluster formation (transition from isolated cell to cell cluster) | Day 1 vs Day 7 | VGP, 2 mg/ml C | Similar | Similar | Similar |
|  | Day 1 vs Day 7 | MET, 2 mg/ml C | Increase | Similar | Decrease |
|  | Day 1 vs Day 7 | VGP, 4 mg/ml C | Increase | Decrease | Increase |
|  | Day 1 vs Day 7 | MET, 4 mg/ml C | Increase | Increase | Increase |
