## Supplement Table S2 Experimental design for "Intracellular mechanics and TBX3 expression jointly dictate the spreading mode of melanoma cells in 3D environments"

Table S2. Summary of experimental design and time points of assessments

| **Activity, assessment** | **Time point (Day)** | | | | |
| --- | --- | --- | --- | --- | --- |
|  | **0** | **1** | **4** | **7** | **10** |
| Cell embedding | X |  |  |  |  |
| Morphology |  | X | X | X | X |
| Microrheology |  | X |  | X |  |
| TBX3 |  | X |  | X |  |
| Actin |  |  |  | X |  |
